# Spatial probabilistic patterns of joint heat and water stress of chickpea in Australia

**DOI:** 10.1101/2025.07.09.664009

**Authors:** Lachlan Lake, James B. Manson, Alan D. Severini, Yash Chauhan, Karine Chenu, Millicent R. Smith, Victor O. Sadras

## Abstract

Quantitative characterisation of the crop environment for breeding and agronomic applications is commonly based on the cluster analysis of isolated climate factors, such as drought and temperature. The focus on individual climate factors fails to account for both the correlations between climate factors in space and time, and the contemporary models of dynamic biological systems where multiple interacting factors simultaneously shape the crop phenotype. With a focus on chickpea in Australia, our aims are to (1) assess the associations between climate factors and actual yield measured in 578 location -years; associations are investigated on a biologically relevant developmental scale; (2) establish the spatial, probabilistic patterns of multivariate environment types at regional and continental scales; and (3) evaluate the shifts in the frequency of environment types with climate change.

We identified a syndrome of hot, dry and high vapour pressure deficit with three intensities of stress classified as low, medium and high. Measured yield, in the range from failed crop to 4 t ha^-1^, aligned more strongly with this multivariate syndrome than with isolated climate factors, with stronger associations at regional than continental scale. The frequency of stressful environment types increased with realised climate change and was most severe in the chickpea growing heartland of central Queensland; our models show that increased frequency of stress could be partially mitigated with earlier flowering varieties. Our findings will inform chickpea agronomy, phenotyping and breeding.

## 1 Introduction

Farmers’ more important economic decisions are made at the business and farm scales, including the allocation of land, capital, labour, and other resources among the alternatives (Kirkegaard, 2019; Rodriguez & Sadras, 2011). Once the farm scale decisions are made, farmers use two primary technologies for sustainable production: varieties and agronomy, i.e., what to grow and how. On historic scales, breeding and agronomy have contributed almost equally to increasing crop production (Bell et al., 1995; Fischer, 2009; Liu et al., 2021; Slafer & Andrade, 1989). For rainfed crops, pairing variety and sowing date is critical to manage the trade-off between fast development and frost risk, and slow development and risk of heat and drought (Collins & Chenu, 2021; Flohr et al., 2017; Lake et al., 2021; Zheng et al., 2018).

Crop adaptation to climate stress including frost, heat and drought, depend on the timing, intensity, and duration of the stress event in relation to crop phenological stage (Darwin, 1859; Prasad & Djanaguiraman, 2014). In addition, the phenotype of stressed crops vary with conditions before the stress event through acclimation and after the event through compensatory mechanisms (Alberdi et al., 1993; Berenguer & Faci, 2001; Campbell et al., 2007; Jifon & Wolfe, 2005; Lee et al., 2005; Marcellos & Single, 1984). Hence, quantifying the major stress patterns is relevant to both breeding and agronomy in a context of risk management.

Jordan and Miller (1980), drew hypothetical patterns of soil moisture stress on a phenological scale to make explicit that phenotypes are needed to match stress patterns; in the terms of Tardieu (2012) “any trait or trait-related allele can confer drought tolerance: just design the right drought scenario”. Two decades after Jordan and Miller (1980), Chapman et al. (2000; 2000) formalised a 3-step method for drought characterisation. First, daily water supply:demand ratio is modelled on a phenological scale for the sites and climate series of interest; crop modelling allows simulation of thousands of patterns that can be representative of the target population of environments for agronomists and breeders (Chenu, 2015; Chenu et al., 2011; Chenu et al., 2013). Second, cluster analysis is used to consolidate a handful of agronomically meaningful stress types. The agronomic relevance is further tested through the alignment of stress types and yield. Often this step involves modelled yield, but independent yield data avoids circularity (Chenu et al., 2011; Elmerich et al., 2023; L. Lake et al., 2016; Manson et al., 2025; Sadras et al., 2012b). Third, the frequency of each stress type is mapped to return a spatial, probabilistic account of drought. A similar approach has been used to characterise thermal regimes in pulses (Lachlan Lake et al., 2016; Sadras et al., 2012a) and nitrogen stress in wheat (Ly et al., 2017). To account for the lack of independence between stresses, e.g. drought and heat often correlate, a series of approaches have been advanced to combine stress patterns (Beillouin et al., 2018; Couëdel et al., 2021; Elmerich et al., 2023); Manson et al. (2025) critically compares these approaches.

For chickpea in Australia, four types of drought stress and three thermal regimes for maximum and minimum temperature have been characterised individually on a continental, but not regional scale (L. Lake et al., 2016). Regional scaling are important in Australia where latitude and continentality are large sources of variation in climate (Gentilli, 1971). In addition to daylength, mean temperature and vernalisation, soil moisture modulates phenological development in chickpea (Li et al., 2022). Lack of consideration of soil moisture impact on chickpea phenology may have biased stress patterns in a previous environmental characterisation (Lachlan Lake et al., 2016). Algorithms that account for soil moisture improve the prediction of flowering and podding and shift the trade-off between stresses whereby earlier flowering in dry soil increases frost risk and reduces heat and drought stress (Chauhan et al., 2019; Li et al., 2022). Accordingly, earlier flowering in dry soil increases frost risk and reduces the risk posed by heat and drought in the critical podding stage (Lake & Sadras, 2014).

Shifts in the patterns of environmental stress with climate change have not been investigated in chickpea. Historically, low temperature has been assumed to delay podding in flowering crops, but a multivariate perspective supports a syndrome of high humidity, and low temperature and radiation that delays podding with implications for harvest index and yield (Gimenez et al., 2024). These findings add to the evidence of plants sensing and responding to the multivariate properties of the environment (Aphalo & Sadras, 2022; Casal & Balasubramanian, 2019).

In the context of industry investments to improve stress adaptation in chickpea, we use an updated phenology model that captures the effect of soil moisture (Chauhan et al., 2024) to address three aims. First, to assess the associations between environmental factors and actual yield measured in 578 site-seasons; associations are investigated on a biologically relevant developmental scale. Second, to characterise the nature and the frequency of occurrence of univariate and multivariate environment types at regional and continental scales for the most impactful environmental factors identified in aim 1. Third, to evaluate the shifts in the frequency of environment types with climate change.

## 2 Method

### 2.1 Overview

Two studies were conducted: Study 1 to identify impactful environmental factors (Aim 1) and Study 2 to characterise the nature and spatial variability of environment types (Aim 2) and evaluate how they have shifted over time (Aim 3). In both studies we simulated crop phenology and water supply : demand ratio using the chickpea module in the Agricultural Production Systems sIMulator, APSIM version 7.10 (Holzworth et al., 2014). This model has been tested across Australian environments for phenology and yield, and includes an algorithm accounting for soil moisture to modulate phenological development that is not available in other versions of the model (Chauhan et al., 2024). The model was run with climate data from SILO on the Queensland Government, Long Paddock website (www.longpaddock.qld.gov.au/silo/) and soil data from the APSoil data base (https://www.apsim.info/apsim-model/apsoil/). Sowing date was actual (Study 1) or based on a sowing rule with a minimum of 15 mm over 5 days (Study 2). In both studies, soil water was reset to the median soil moisture from each studied site on 1-Jan assuming a wheat-chickpea rotation. The median was determined by running continuous wheat from 1957 to 2023 and retrieving the simulated soil water on Jan 1. Manson et al. (2025) justify this approach to initialise soil moisture.

In both studies, we characterised environments based on a daily timestep for maximum and minimum temperature, heat stress and frost frequency, rainfall, water supply:demand, vapour pressure deficit and radiation. The water supply:demand ranges from 1, no stress to 0, maximum stress (Chapman 2000ab). We defined thresholds for heat stress, i.e., maximum temperature > 30°C (Devasirvatham et al., 2015; Jeffrey et al., 2021), chilling temperature compromising pod set, i.e. mean temperature < 15°C (Croser et al., 2003), and frost, i.e., minimum temperature < 0 °C (Croser et al., 2003). Crop development was centred at the simulated time of flowering and the season divided into sequential 200 °Cd-windows from -1000 °Cd before flowering to 1200 °Cd after flowering; for the water supply:demand cluster analysis, the scale was truncated at 800 °Cd after flowering because modelling of canopy senescence and maturity are unreliable (Manson et al., 2025). However, the average values are still presented for completeness. Methods specific to each study are in the next sections.

### 2.2 Aim 1: associations between environmental factors and actual yield measured in 578 trials

We used measured yield from an extensive rainfed National Variety Trial (NVT, <u>nvt.grdc.com.au</u>) data set including 27 Northern, 18 Southern and 21 Western growing environments spanning 19 years (2005-2023), returning 578 location-year combinations (Fig. 1). The NVT database includes yield for desi and/or kabuli chickpea, but we used the data for desi for three reasons: (i) desi was represented in all location-years, (ii) the pairwise comparison between desi and kabuli yield aligned with the 1:1 line (Fig. 2a), and (iii) the yield difference between kabuli and desi was small compared with the variation between location-years from failed crop to 4 t ha^-1^ (Fig. 2b-e). Using actual sowing date for each location-year, we simulated phenology and water supply:demand ratio using cultivar parameters for Sonali (early phenology) and PBA HatTrick (standard phenology). Next, we calculated Pearson’s correlation coefficient and its *p*-value for the associations of measured yield with environmental factors in each 200 °Cd-window. We controlled the false discovery rate of the *p*-values with the Benjamini-Hochberg method (Benjamini & Hochberg, 1995). After initial analysis we decided to centre the weather data on flowering time of PBA HatTrick because its phenology represents standard varieties in all three regions, and the correlation coefficients were hardly affected by the choice (Sup. Fig. 1).

**Figure 1.**
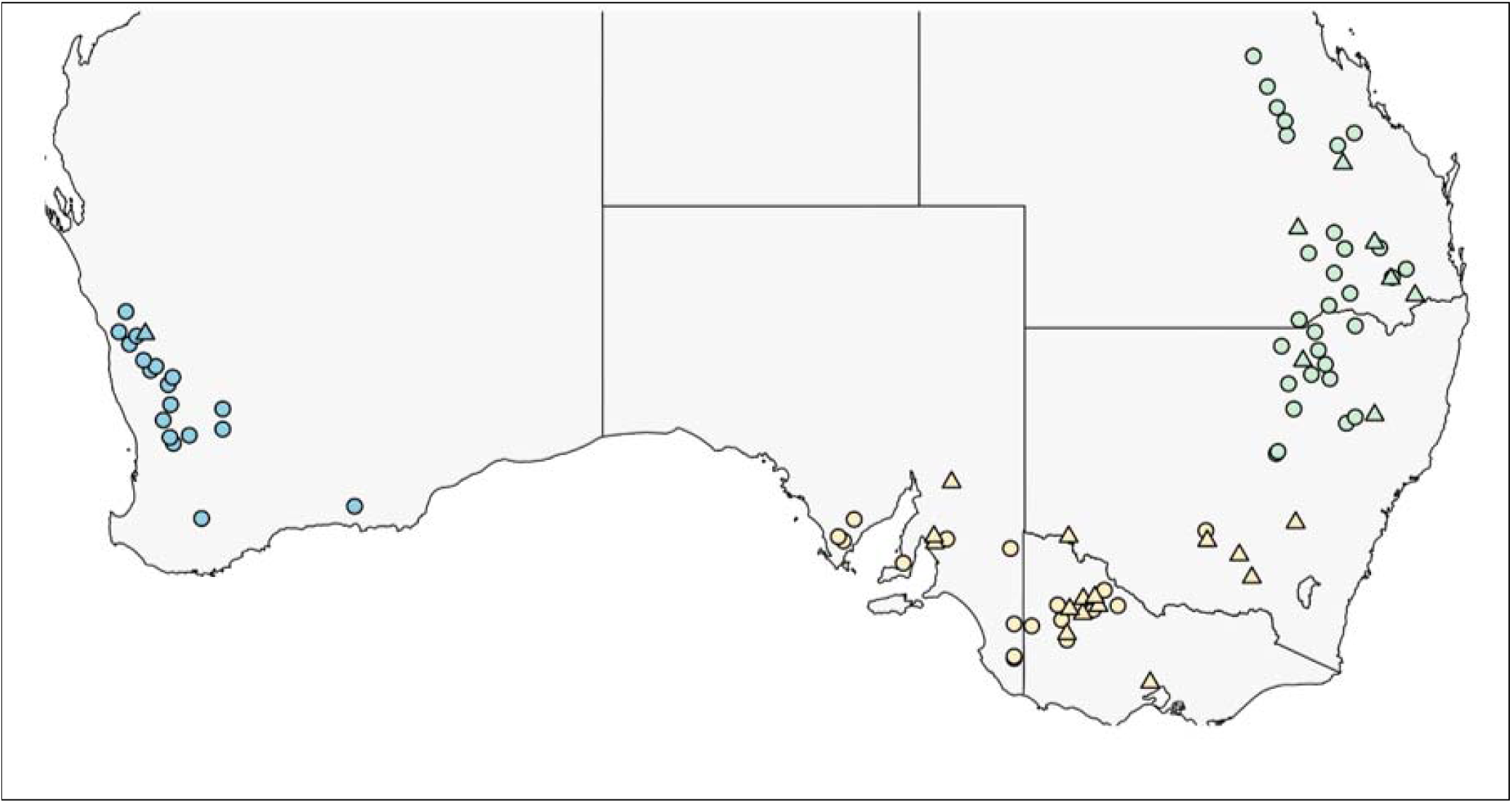
National Variety Trials (circles) and Chickpea Breeding Australia trials (triangles) used for the selection of locations in the study. Colours show Northern (green), Southern (yellow) and Western (blue) growing regions.

**Figure 2.**
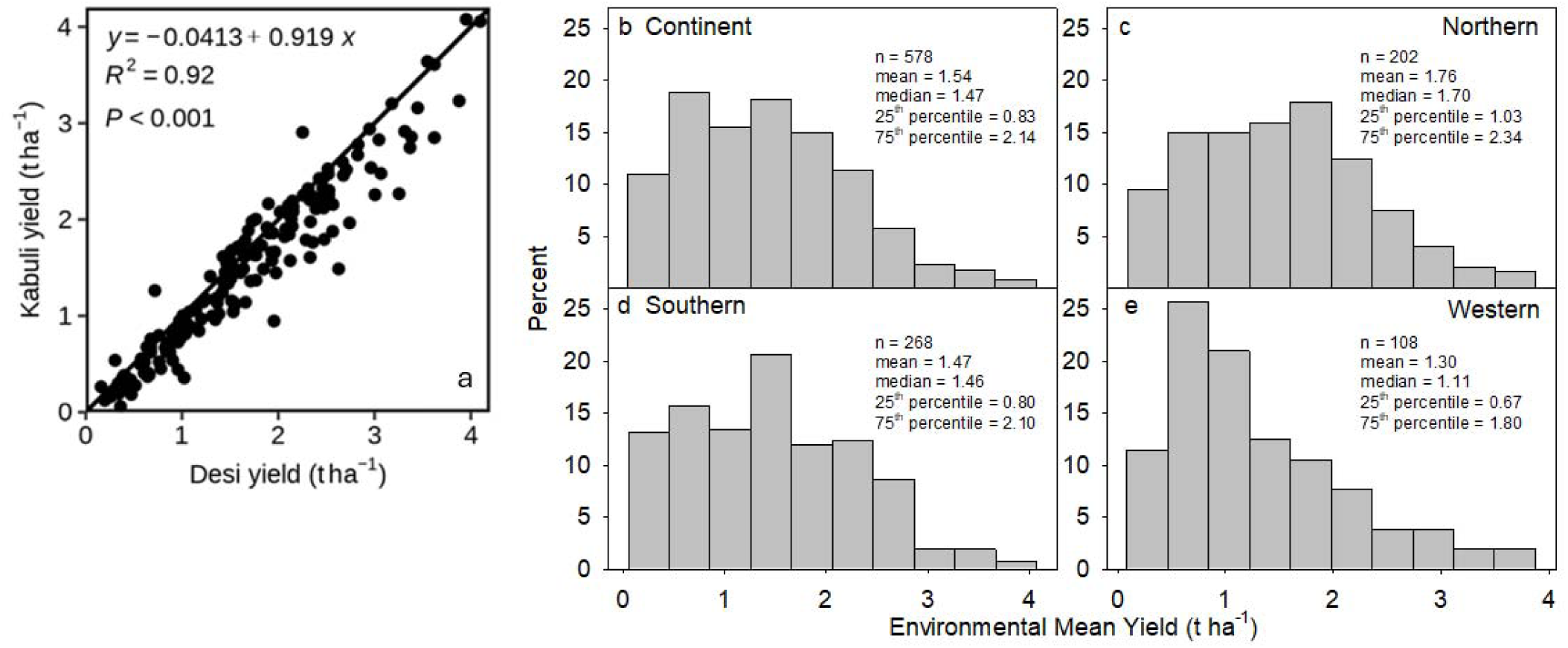
Frequency distribution of trial mean yield measured in kabuli and desi varieties in 578 National Variety Trials between 2005 and 2013, (A) nationally and (B-D) regionaly. (E) Correlation between yield of kabuli and desi varieties; the line is y = x.

To determine correlations between environment types (determined as in section 2.3) and measured NVT yield, we conducted a separate cluster analysis; environmental data for NVT location-years were joined to the larger environmental characterisation dataset (NVT and Chickpea Breeding Australia (CBA) locations, 1957 to 2023, 2 varieties) and clustered so that we could match NVT yields to environment types. It is not likely that the addition of a relatively small number of location-years changes the environment typing outcome, but we checked to confirm that there was no effect on environment types for each analysis. For each environment type, we calculated the 10-25-50-75-90th percentiles of measured yield.

### 2.3 Aim 2: characterisation of the nature and frequency of univariate and multivariate environment types

To quantify the major chickpea environment types, we chose 93 locations that captured the spread of the national growing area (Fig. 1). We performed cluster analysis using simulations and climate data for the factorial combination of 93 locations, 66 years (1957-2023), and two varieties, Sonali and PBA HatTrick. Cluster analysis was conducted for each environmental factor at (i) the regional and continental scales, and (ii) individually and for a combination of factors using the method in Manson et al. (2025). Briefly, we used the CLARA (“Clustering LARge Applications”) algorithm to cluster environmental factors (Kaufman & Rousseeuw, 2009). This was performed with the clara() function from the cluster package in the R environment (R Development Core, 2014), using the Euclidean distance measure on standardised data (mean = 0, standard deviation = 1) and the optimal number of clusters as identified from the knee/elbow method for average silhouette width, total dissimilarity, and R^2^. For water supply:demand ratio, maximum temperature and VPD for all locations and years from 1957 to 2023, we performed the cluster analysis on 200 °Cd windows from - 1000 °Cd before flowering to 1200 °Cd after flowering (except supply demand which was truncated, see above), at regional and continental scales. We limited our characterisations to these variables because they had the strongest correlations with yield (Aim 1). We ranked the univariate and multivariate environment types (ETs) from the least stressful ET1 to the most stressful ET3; we assigned the severity ranking of the stress environment types based on both their biological fit and yield ranking.

To address the spatial scaling, the patterns of regions and continent were compared visually and analytically. For each location-year, we compared their regional ET to their continental ET. We then calculated the frequencies that location-years had (i) the same rank ET rank, (ii) ranks that differed by 1 (e.g. ET1 vs ET2) and (iii) ranks that differed by 2 between regional and continental scales. Similarly, comparisons were made to investigate interchangeability between single and multivariate environment types. A high frequency of location-years assigned to the same or similar ranks indicates a high level of agreement between the clustering results and therefore a high level of scalability, region and continent, or interchangeability, multivariate and univariate, of the characterisations.

### 2.4 Aim 3: shift in environment types with climate change

Our environmental characterisations use a phenological scale for biological significance. Hence, to investigate shifts in environment types with climate change, we first compared simulated phenology using actual weather data and modelled phenology using simulated climate projections with model Access CM2 (https://research.csiro.au/access/about/cm2/), a model previously downscaled to a resolution of 10 km using the Conformal Cubic Atmospheric model (CCAM, (Chapman et al., 2023)) for an overlapping period from 2015 to 2023. These gridded weather data were processed to extract daily values from the specific coordinates of each site using linear interpolation with the software CDO (Schulzweida, 2021). The comparisons showed a mismatch of up two weeks in phenology predictions using actual and simulated weather. This was primarily associated with unrealistic projections of rainfall patterns that biased the dry-wet sequences. For example, in Billa Billa (−28.15, 150.25), actual number of rainy days per year was 91 compared to 321 simulated with Access CM2. Owing to lack of reliable daily-scale projections in rainfall that is required to model phenology with APSIM accounting for soil moisture, we compared the frequency of environment types in the standard three-decade periods of the World Meteorological Organisation for ‘present’ climate, 1991-2023, and ‘historic’ climate, 1961 to 1990 (Hulme, 2020).

## 3 Results

### 3.1 Aim 1: associations between environmental factors and actual yield

Figure 2 summarises actual yield data in 578 location -years. The yield of desi and kabuli aligned across environments (Fig. 2a) and is thus pooled in the histograms of yield distributions at the regional and continental scale. Nationally yield varied from failed crop to 4 t ha^-1^ and averaged 1.5 t ha^-1^. Median yield was 1.10 t ha^-1^ in the Western, 1.46 t ha^-1^ in the Southern and 1.70 t ha^-1^ in the Northern region (Fig. 2c-e).

Figure 3 shows the associations between environmantal factors and measured yield on a phenological scale. Yield associated strongly with maximum temperature, heat stress frequency, and water stress, with r exceeding 0.5 for water supply:demand ratio. There was no association with frost; frost is highly dependent on topography and was not well captured in our interpolated weather dataset (Manson et al. 2025).

**Figure 3.**
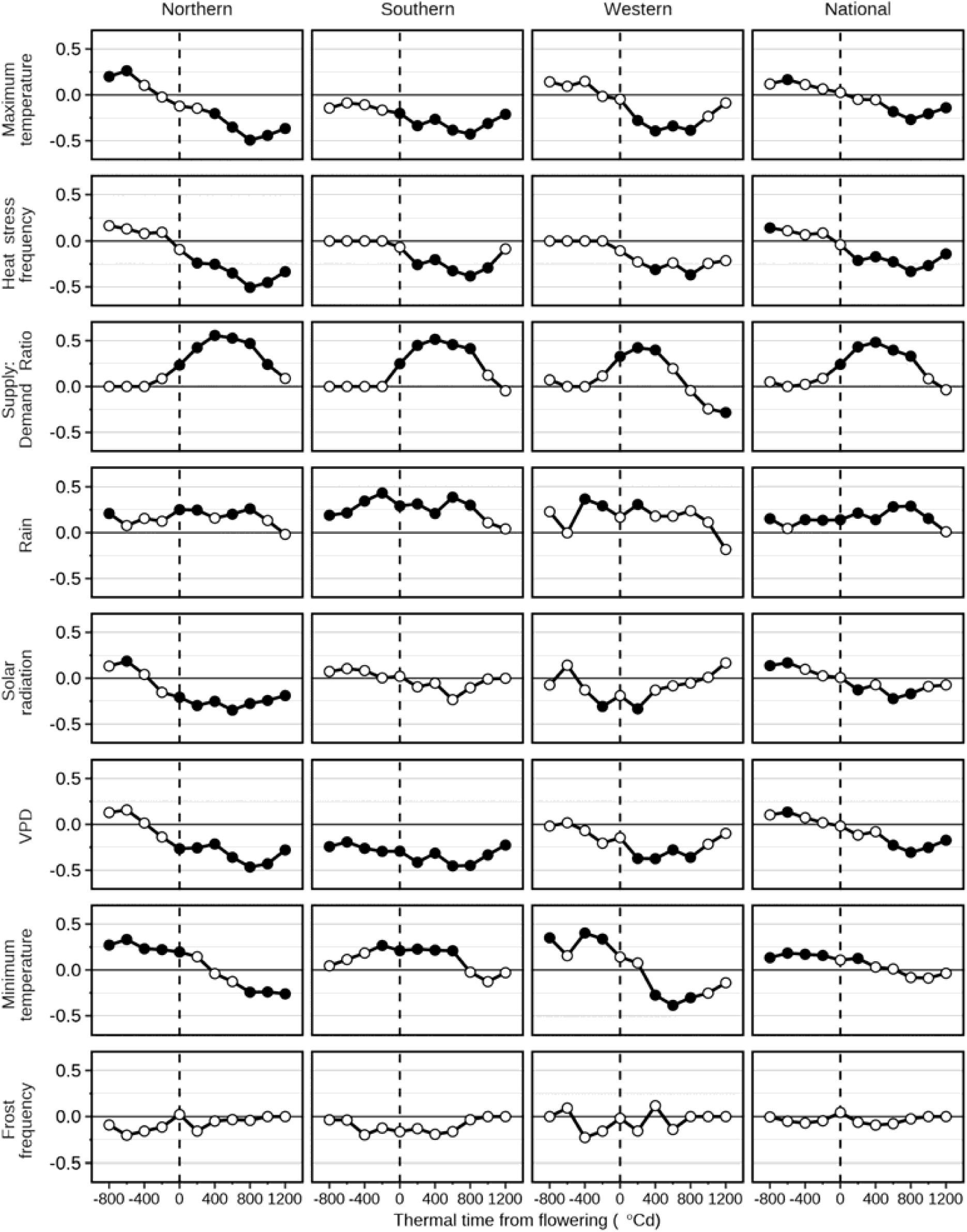
Pearson’s coefficient of correlation between Nataional Variety Trial mean yield and eight environmental factors. Factors are centred on the average simulated flowering date, and averaged every 200 °Cd. Correlations presented at the regional and continental scales. Symbols indicate Benjamini-Hochberg adjusted *p* ≤ 0.05 (closed) and *p* > 0.05 (open). We defined thresholds as heat stress, is maximum temperature > 30°C, and frost, i.e., minimum temperature < 0 °C. VPD is vapour pressure deficit. Phenology and water supply:demand were modelled with genotypic parameters for PBA HatTrick.

Maximum temperature had a consistent negative association with yield after flowering across all three regions and a positive correlation before flowering in the Northern region; heat stress frequency followed the same pattern. As they are linked, VPD followed a pattern similar to both maximum temperature and heat stress frequency. Water stress showed a negative relationship with yield beginning at flowering across regions. The Western region was unique with shorter duration of the correlation and with dry conditions at the end of the growth period favouring yield. Rainfall followed a similar trend to water supply:demand but with weaker correlations with yield both regionally and continentaly. In the Northern and Western regions, minimum temperature was positively associated with yield before flowering and negatively after flowering, whilst in the Southern region it was associated with yield before and after flowering.

### 3.2 Aim 2: univariate and multivariate environment types

Environment types were characterised for for the three variables with the largest associations with yield, i.e. maximum temperature, water supply:demand ratio and VPD, as both univariate (Fig. 4) and multivariate environmental types (Fig. 5). Environment types for maximum temperature and VPD showed greater geographical differences compared to water supply:demand. For example, the least stressful environment types (ET1) of maximum temperature and VPD in the Northern region were similar to the most stressful types of the Southern region; continental environment types were close to the average of all three regions.

**Figure 4.**
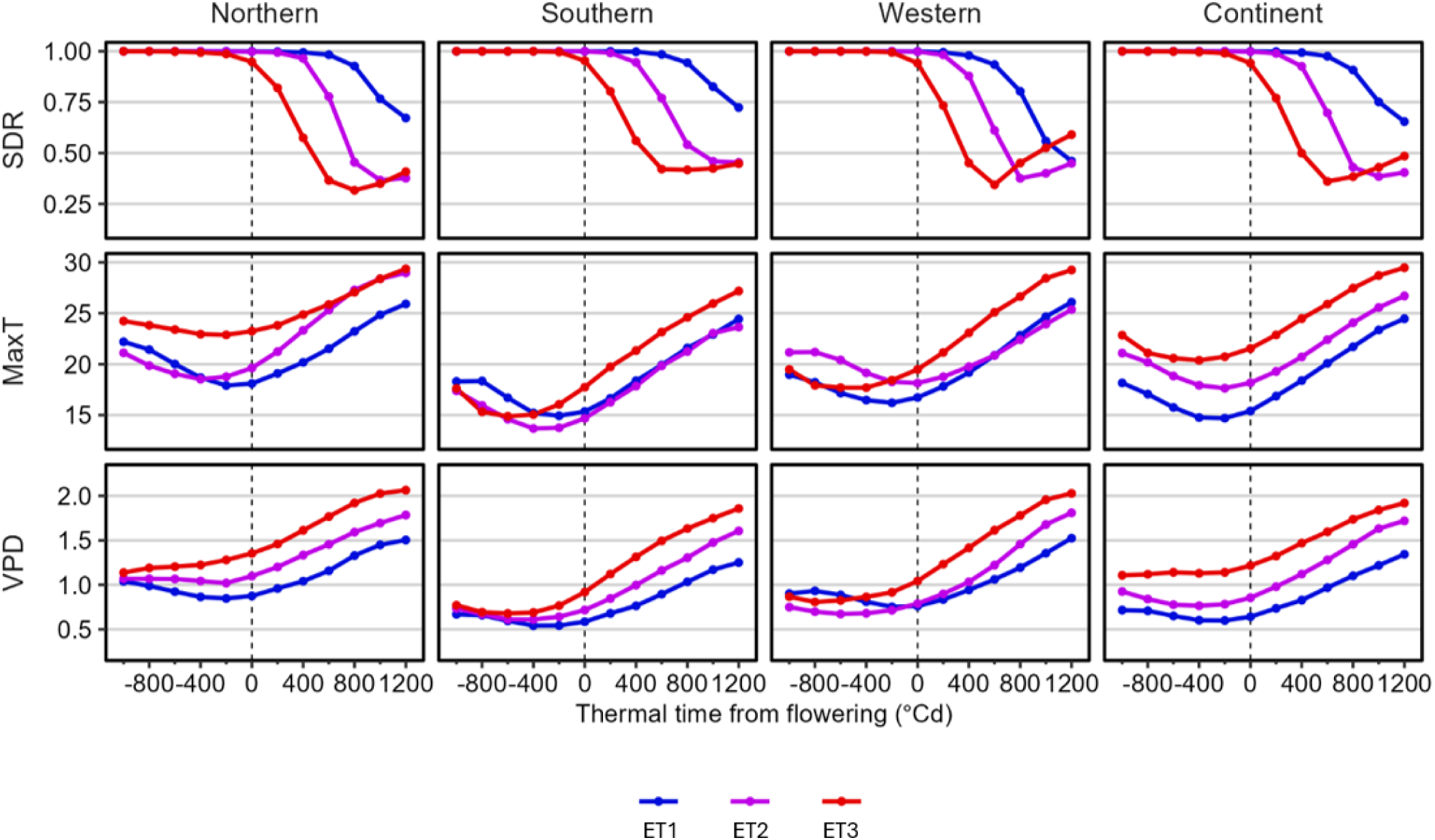
Univariate environment types of water supply:demand ratio (SDR), maximum temperature (MaxT) and vapour pressure deficit (VPD), for Northern, Southern, Western regions and for the continent. Points are the average of each cluster, blue is the least stressful (ET1), red is most stressful (ET3), and purple is intermediate (ET2). Environmental data are from simulations for two chickpea varieties (early, late) across 93 locations across Australia from 1957 to 2023. The SDR cluster analysis was performed on truncated data up to 800 °Cd after flowering, however trends are presented with average data up to 1200 °Cd for completeness.

**Figure 5.**
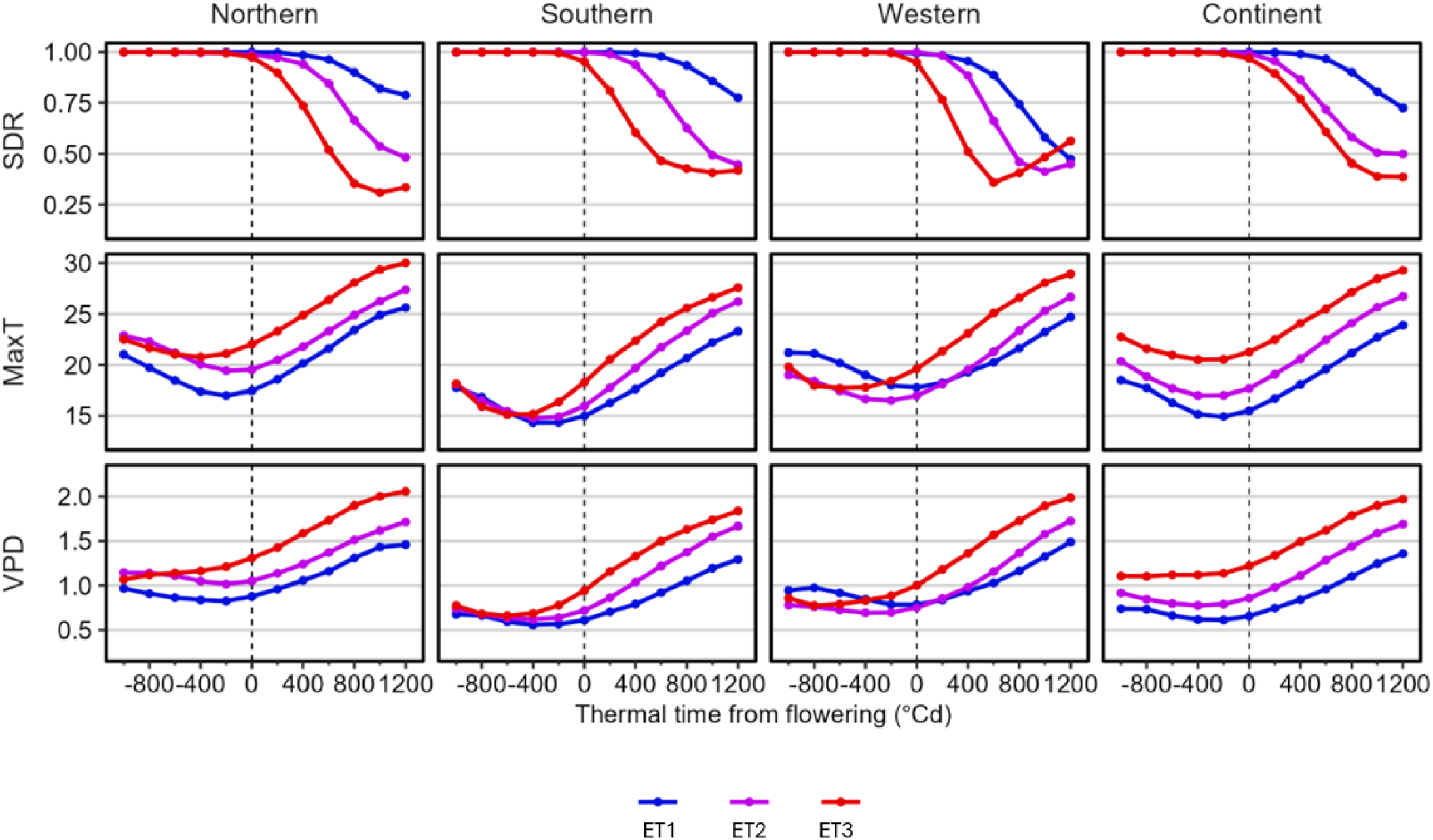
Multivariate environment types of water supply:demand ratio (SDR), maximum temperature (MaxT) and vapour pressure deficit (VPD), for Northern, Southern, Western regions and for the whole continent. Points are the average of each cluster, blue is the least stressful (ET1), red is most stressful (ET3), and purple is intermediate (ET2). Environmnetal data are from simulations for two chickpea varieties (early, late) across 93 locations across Australia from 1957 to 2023.

Water supply:demand ratio showed the closest agreement between continental and regional scales: 87 to 91 % of location-years were assigned to the same environment type rank (e.g., location-year with ET1 in one region were mostly ET1 in the other regions and at the contiental scale, Table 1). In all regional and continental water supply:demand environment types, there was almost no stress before flowering even in the most stressful ET3.

**Table 1.** Frequency of agreement in the assignation of location-years to the same rank between regional and continental scales, or differed by 1 (ET1-ET2) or 2 (ET1-ET3) ranks.

| Region | Env.type | Frequency comparing regional ET rank versus continental ET rank |  |  |
| --- | --- | --- | --- | --- |
|  |  | Same rank | Ranks differing by 1 | Ranks differing by 2 |
| Northern | MaxT | 0.42 | 0.58 | 0.00 |
|  | SDR | 0.90 | 0.10 | 0.00 |
|  | VPD | 0.41 | 0.56 | 0.02 |
|  | Multivariate | 0.63 | 0.36 | 0.00 |
| Southern | MaxT | 0.33 | 0.55 | 0.13 |
|  | SDR | 0.91 | 0.09 | 0.00 |
|  | VPD | 0.62 | 0.38 | 0.00 |
|  | Multivariate | 0.74 | 0.26 | 0.00 |
| Western | MaxT | 0.65 | 0.35 | 0.00 |
|  | SDR | 0.87 | 0.13 | 0.00 |
|  | VPD | 0.57 | 0.42 | 0.01 |
|  | Multivariate | 0.60 | 0.40 | 0.00 |

Frequencies for the water supply:demand ETs associated with regional rainfall seasonality (Sup. Fig. 2). Regionally and continentally the most stressful environment type (ET3) was the least frequent in the summer rainfall environments of the north, but the most common in various locations in the winter-rainfall environments of the west and south. In the Southern region, the dominant ET was the least stressful.

Maximum temperature showed a partial disagreement between continental and regional scales: 33 to 65 % of location-years were assigned to the same ET rank (Table 1). At a continental scale, the most stressful environment was hotter for the entire season and the least stressful being coolest; regional patterns were distincly different. At the start of the year in the Northern region, the most stressful environment was the hottest and the medium stress environment was the coolest, but 400 °Cd after flowering the positions reversed with the most stressful conditions becoming cooler than the medium conditions. Conversely, in the Southern region, the most stressful conditions began the season coolest, becoming hottest 400 °Cd before flowering onwards. Frequencies for the environment tyes differed between regions with the hottest and most stressful ET3 being the most frequent in the north and Western, but medium stress ET2 being most frequent in the Southern region. Least stressful conditions were ubiquitiously least frequent.

Vapour pressure deficit was similar to maximum temperature and showed a partial disagreement between continent and regional scales: 41 to 57 % of location-years were assigned to the same environemt type rank (Table 1). The most and least stressful environment types for the Northern, Southern and continental scales showed corresponding higher and lower VPD across the season. However in the Western region, the least stressful environment type ET3 began with the highest VPD and finished with the lowest.

The multivariate environment types (Fig. 5) were partway between maximum temperature and water supply:demand with 60 to 74 % agreement in rankings (Table 1). Across all comparisons, almost no location-years were assigned to opposite environment types (ET1 and ET3) between continent and regions, i.e. 0 to 2 % differed by two ranks or most and least stressful. The exception was environment types for maximum temperature in the Southern region where the mistmatch was 13%.

#### 3.2.1 Spatial scaling of patterns and scaling from single to multivariate characterisations

Table 2 shows the agreement between ranking of single and multivariate environment types determined at the continental and regional scales. Rankings at the continental scale show that agreement in environment types was 80% for maximum temperature, 87% for VPD and 56% for water supply:demand. The frequency of mistmatch in environmental types defined at regional scale was 45 - 66% for maximum temperaturem, 21 - 41% water supply:demand and 21 - 25% for VPD.

**Table 2.**
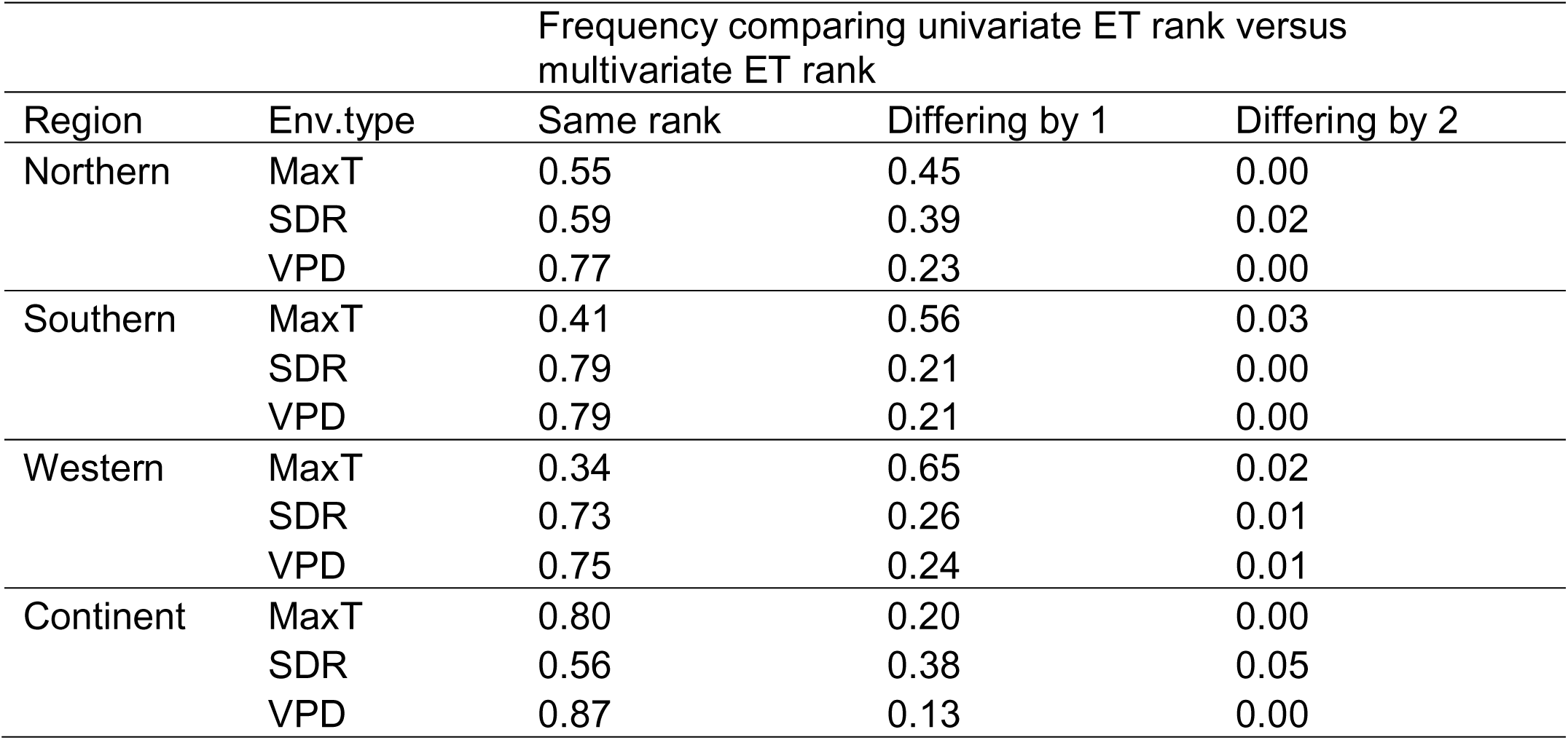
Frequency of agreement in the assignation of location-years to the same rank between multivariate and univariate components ET, or differed by 1 (ET1-ET2) or 2 (ET1-ET3) ranks. Comparison were performed for the maximum temperature (MaxT), water supply:demand ratio (SDR), and vapour pressure deficit (VPD) comparing regional ET together, and continental ET together.

| Region | Env.type | Frequency comparing univariate ET rank versus multivariate ET rank |  |  |
| --- | --- | --- | --- | --- |
|  |  | Same rank | Differing by 1 | Differing by 2 |
| Northern | MaxT | 0.55 | 0.45 | 0.00 |
|  | SDR | 0.59 | 0.39 | 0.02 |
|  | VPD | 0.77 | 0.23 | 0.00 |
| Southern | MaxT | 0.41 | 0.56 | 0.03 |
|  | SDR | 0.79 | 0.21 | 0.00 |
|  | VPD | 0.79 | 0.21 | 0.00 |
| Western | MaxT | 0.34 | 0.65 | 0.02 |
|  | SDR | 0.73 | 0.26 | 0.01 |
|  | VPD | 0.75 | 0.24 | 0.01 |
| Continent | MaxT | 0.80 | 0.20 | 0.00 |
|  | SDR | 0.56 | 0.38 | 0.05 |
|  | VPD | 0.87 | 0.13 | 0.00 |

We present the continental correlations between environmental factors between 200 °Cd to 600 °Cd after flowering, i.e. the period when environmental variables had the largest variation and strongest correlations with actual yield, in Table 3. Regional sclaes were similar and presented in Supplementary Tables S1 to S3. Water supply:demand was strongly negatively correlated to maximum temperature, radiation and VPD, while maximum temperature was strongly correlated with radiation and VPD.

**Table 3.** Correlation coefficients for the average water supply:demand ratio (SDR), maximum temperature (MaxT), minimum temperature (MinT), solar radiation (Radn) and vapour pressure deficit (VPD) from 200 °Cd to 600 °Cd after flowering across the 6138 location-years studied.

|  | Supply:demand | Maximum Temperature | Minimum Temperature | Radiation |
| --- | --- | --- | --- | --- |
| Maximum Temperature | -0.52 |  |  |  |
| Minimum Temperature | -0.20 | 0.46 |  |  |
| Radiation | -0.47 | 0.65 | 0.07 |  |
| VPD | -0.54 | 0.96 | 0.20 | 0.70 |

#### 3.2.2 Agronomic significance of environment types

To assess the agronomic significance of the simulated environment types, we explored their alignment with measured NVT yield (Study 1, Fig. 2) (Fig. 6). On a continental scale, the environment type for water supply:demand ratio followed by the multivariate environment type achieved the greatest discrimination of observed yield. Maximum temperature and VPD environment types showed little discrimination for yield; both showed similar yield trends for the low and high stress environment types, with medium stress having the lower yield.

**Figure 6.**
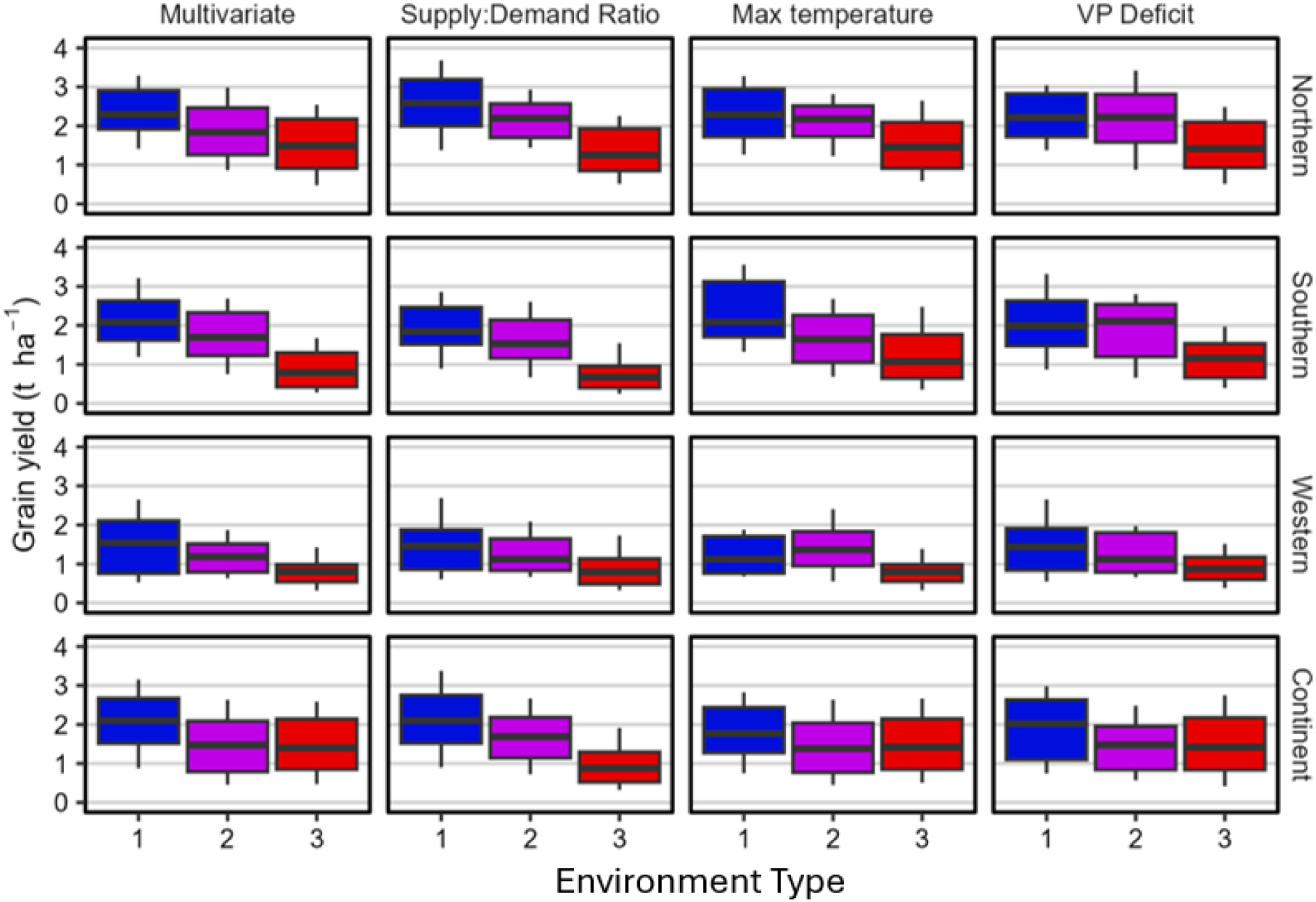
Frequency distribution of National Variety Trial mean yield in environment types for multivariate and individual environmental factors of water supply:demand ratio, maximum temperature and vapour pressure deficit in each region and the whole continent. Boxes are 25-50-75th percentile, tails are 10-90th percentile. Stress intensity increases from 1 to 3 (ET1-ET3) and from blue to purple to red. The environment types are presented in Figs 4 and 5.

On a regional scale, the multivariate environment types featured superior discrimination of yield (Fig. 6). Water supply:demand ratio had less resolution to discriminate between low and medium stress in the Western region, and to a lesser degree in the Southern. Maximum temperature environment types aligned with yield in the Southern region but not in the Northern or Western regions. Yield did not align with VPD environment types. The poor alignment of yield and VPD environment types may reflect the confounding correlations with water supply:demand, maximum temperature, and radiation (Table 3).

#### 3.2.3 Spatial variation in frequency of environment types

The frequency of multivariate environment types varied spatially (Fig. 7). For the continental environment types, ET3, which is a syndrome of high water stress, temperature and VPD, rarely occured in southerly latitudes. Conversely ET1, a syndrome of wet and cool conditions rarely occured in Northern latitudes. The lack of differentiation between multivariate environment types at the continental scale suggests regional envrionment types are required for agronomically useful characterisations.

**Figure 7.**
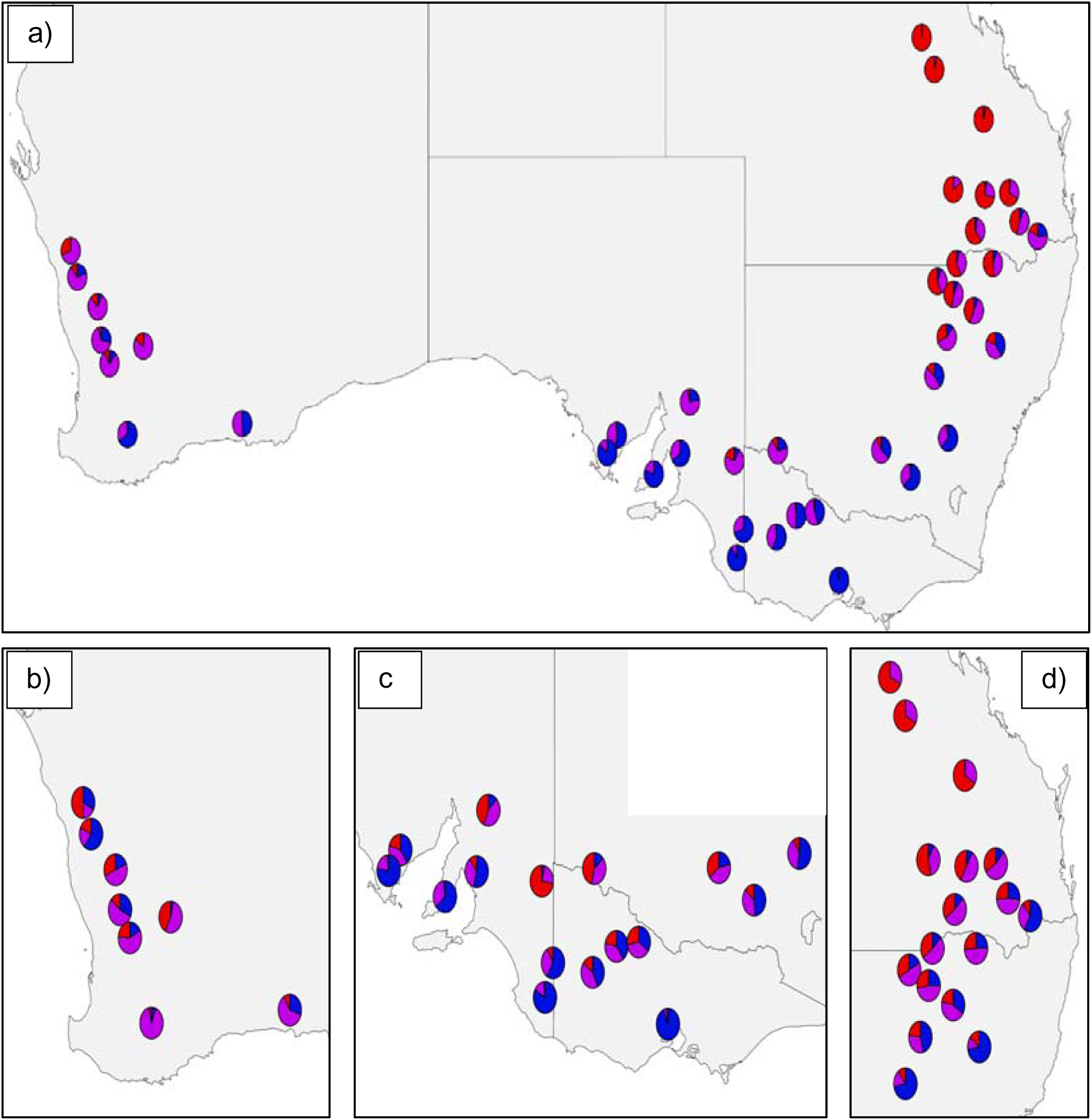
Spatial variation in the frequency of multivariate environment types combining water stress, maximum temperature and VPD for a) environment types identified for the whole continent and environment types identified in each of the b) Western, c) Southern and d) Northern regions. Nearby locations with similar frequency profiles were grouped to minimise overlap between pie charts. Based on simulations from 1957 to 2023. The characteristics of ETs at the regional and continental scales are presented in Fig. 5.

### 3.3 Aim 3: shift in environment types with climate change

Realised climate change has led to the increase of the most stressful environment types in many locations, but there was spatial variaton (Fig. 8). At the continental scale, realised climate change has been more severe in Southern Queensland and Northern New South Wales where multivariate ET3 was already highly frequent. However, ET3 had a relatively smaller increase than in regions where it was historically less frequent. Analysis at a regional scale provides a clearer picture with greater evidence of realised climate change in Southern latitudes leading to higher frequency of the most stressful ET3 (Fig. 8b-d).

**Figure 8.**
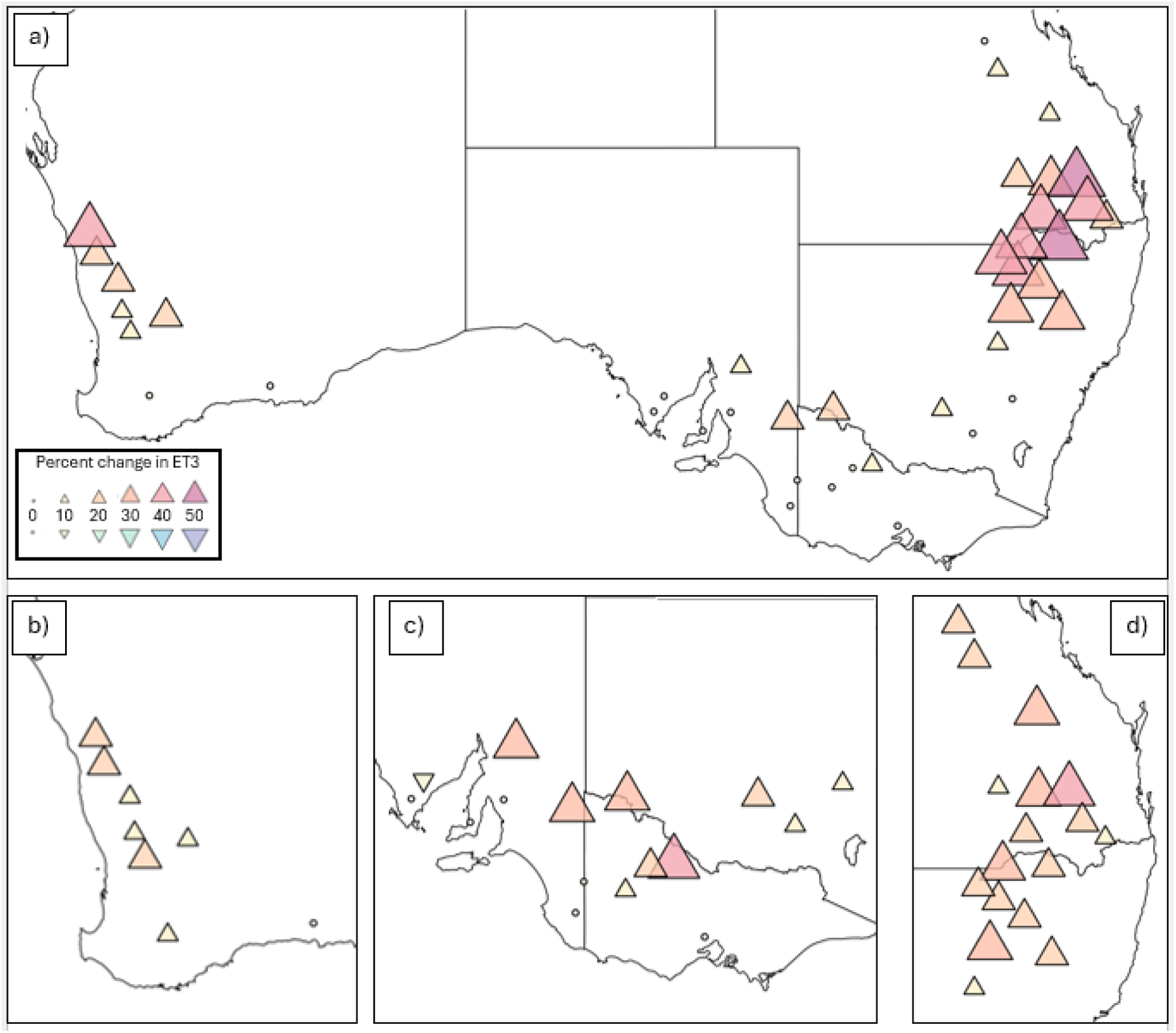
Increase in proportion of the most stressful multivariate environment type (ET3) due to realized climate change (% ET3 under ‘present’ climate (1991-2023) minus % ET3 under ‘historic climate’ (1961-1990)) for a) environment types identified for the whole continent and environmen types identified for the b) Western, c) Southern and d) Northern regions. Nearby locations with similar frequency profiles were grouped to minimise overlap between symbols. The characteristics of ET3s at the regional and continental scales are presented in Fig. 5.

With present climate (1991 to 2023) earlier flowering reduced the frequency of the most stressful multivariate environment type ET3 (Fig. 9). As with the realised climate change (Fig. 8), scaling from continent to region was more informative with the Southern and Northern regions reducing their frequency of multivariate stress (ET3) consistently with earlier flowering.

**Figure 9.**
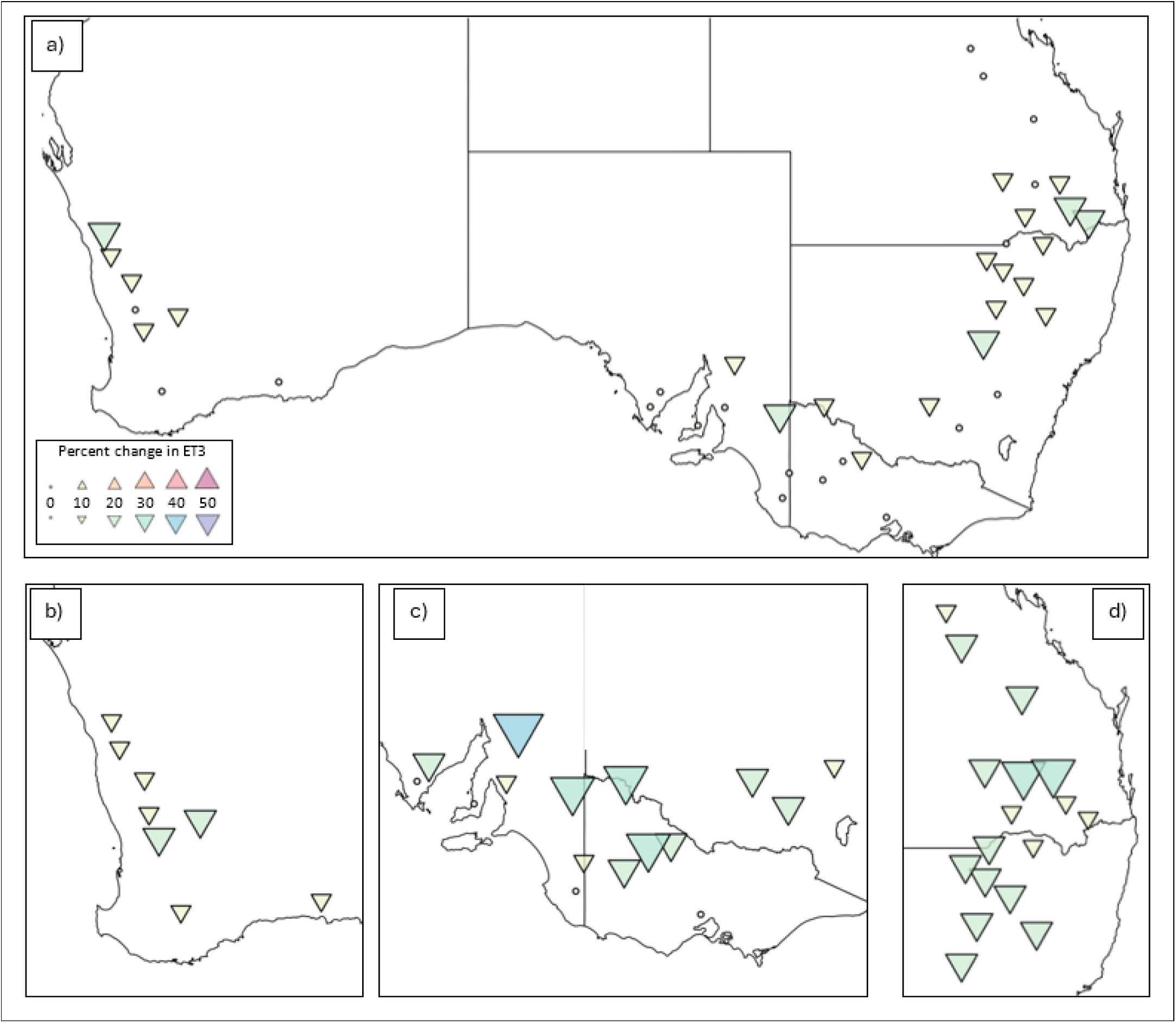
Reduction in proportion of the most stressful multivariate environment type (ET3) due to sowing of an early rather than late maturity type variety (% ET3 with early flowering cultivar minus % ET3 with late flowering cultivar) under present climate (1991 to 2023) for a) the environment type identified for the whole continent and environment type identified for each of the b) Western, c) Southern and d) Northern regions. Nearby locations with similar frequency profiles were grouped to minimise overlap between symbols. The characteristics of ET3s at the regional and continental scales are presented in Fig. 5.

## 4 Discussion

To feed a growing population with higher yielding crops, researchers and breeders will need to use increasingly sophisticated methods to match phenotype and environment (Collins & Chenu, 2021; Cooper et al., 2023; Ravasi et al., 2020). Here we describe different approaches to quantify Australian chickpea environments, focusing on the interactions and correlations between environmental factors, and building on the method of Manson et al. (2025). Additionally we describe ET clusters on both continental and regional scales and explore how phenological change may mitigate stress.

### 4.1 A heat, drought and evaporative demand syndrome is a better explainer of yield

Pre-breeding and phenotyping often target adaptation to climate stresses such as frost, heat and drought in isolation, overlooking interactions that are relevant in the field (Côté et al., 2016; Stoddard et al., 2006). Genetic progress for complex abiotic stress tolerance has been limited and is lagging behind constitutive traits (Aphalo & Sadras, 2022). From biological and agronomic perspectives, a focus on multivariate properties of the environment is justified for three reasons. First, interactions between stresses such as drought and heat or water and nitrogen are common and biologically consequential (Cossani & Sadras, 2018; Côté et al., 2016; Gaur et al., 2019; Hekneby et al., 2006). Second, an information-based model of the phenotype has been developed that considers plants store information on the multivariate properties of the environment on evoultioinary time scale in the genome, in ecological time in the epigenome, and in real time (Aphalo & Ballare, 1995; Aphalo & Sadras, 2022). Examples of plant predictive processes include the priming or acclimation for maximal photosythesis when leaves are exposed to blue light at sunrise; the blue light stimulates phototropins and increases the rate of stomatal opening (Wang et al., 2020). Phytochrome B plays a dual role in sensing both light spectracl composition and temperature (Casal & Balasubramanian, 2019). Aphalo and Sadras (2022) have proposed a hypothesis whereby plant exposure to ultraviolet light plays a role in the pre-acclimation to drought stress. Third, there are stressful and non-stressful effects of environmental factors on crop growth and development. The duration of the critical period for yield in chickpea is determined by the interactions of temperature, photoperiod and soil water stress (Chauhan et al., 2024; Gimenez et al., 2024; Lake & Sadras, 2014; Li et al., 2022). Hence, the main effect of elevated, non-stressful temperature reducing yield is mediated by the shortening of the critical period (Sadras & Dreccer, 2015). A multivariate environmental characterisation of Australian faba bean showed yield was most strongly associated with a sydnrome of drought, heat and high VPD (Manson et al., 2025).

### 4.2 Improved association of yield with multivariate modelling and updated crop model

Our previous environmental characterisation for Australian chickpea was on a continental scale, examined isolated climate factors and used a crop model that did not account for soil water effect on phenology (Lachlan Lake et al., 2016). It identified three heat, three cold and four water stress environment types, with mixed results in describing yield. Mismatches between stress severity and yield occurred with for both maximum and minimum temperature.

Our new continental characterisation of maximum temperature (Fig. 4) partially aligns with the previous study showing high stress environments as being hotter all season and low stress being cooler. Consistent with the previous study, the medium stress ET for maximum temperature was associated with lower yield (Fig. 6).

In terms of water suppply:demand, our new continental characterisation identified only three clusters. The new patterns resembled the previous ones with no stress before flowering across environment types, with varying degrees of severity developing after flowering.

Despite the relatively similar look to the ET clusters, the new ones derived from a model accounting for the effect of soil water on phenology aligned better with yield (Fig. 6).

### 4.3 Regional resolution is required for more efficient classification of thermal environments

Environmental characterisation for chickpea and faba bean in Australia have previously used a continental scale (Lachlan Lake et al., 2016; Manson et al., 2025). This scale pools large pedo-climatically distinct regions (Freebairn et al., 2006; Isbell, 2002) In both previous characterisations, this approach tended to classify regions by one or two dominant environment types. As seen here and in wheat (Chenu et al., 2013), supply:demand scales from region to continent. However, temperature and evaporative demand did not scale; here as in faba bean, the environment type with highest temperautre stress was dominant in the Northern latitudes, with the trend reversing southwards. This pattern led Manson et al. (2025) to sugest that more locally relevant environment types may be identified if the resolution was increased to regional scale. Our results confirm this assesment with more diverse thermal environment types identified at the regional scale (Fig. 7), and with a stronger association with yield (Fig. 6). This is an especially important distinction with previous work when considering that crops in the Northern region are grown predominantly on stored soil moisture while souhtern and Western regions rely primarily on in-season rainfall (Chenu et al., 2013; Sadras & Rodriguez, 2007).

### 4.4 Adaptation to climate change and limitations to future climate modelling

Australian faba bean and wheat are exposed to increased frequency of stress type environments under realised climate change scenarios where drought and heat are prevalent (Ababaei & Chenu, 2020; Manson et al., 2025; Watson et al., 2017). In environments where frost risk is manageable, early flowering and maturity are adaptive in crops such as chickpea and may become increasingly important breeding targets (Berger et al., 2006; Maphosa et al., 2020). Adjusted sowing time may also be useful to reduce stress as demonstrated for cereals under scenarios of increasing frequency of late season drought and heat stress (Collins & Chenu, 2021).

Here, as for faba bean (Manson et al., 2025), we demonstrate the benefits of earlier flowering phenotypes which show significantly reduced frequency of stress exposure under realised climate change (Fig. 9). Using climate projections, drought stress is predicted to increase for European maize (Harrison et al., 2014), South American beans (Heinemann et al., 2017), Western Australian wheat (Watson et al., 2017) and European soybean (Nendel et al., 2023). We wished to explore the effect of future climates on chickpea production environments, but available climate models returned unrealistic rainfall patterns with frequent, e.g. >300 days per annum, and extremely small rainfall events (<0.01mm), which biased phenological predictions with a crop model accounting for effect of soil water (Chauhan et al., 2024).

## 5 Conclusions

We demonstrate improvements in quantitative characterisation of Australian chickpea-growing environments through use of an updated crop model that accounts for the effects of soil water on phenology, the incorporation of multiple environmental factors into a single multivariate cluster, and the scaling to regional level. Out of all the environmental factors studied, drought is the main stressor discriminating for yield, at both regional and continental scales. The incorporation of multiple environmental factors into a single multivariate cluster allowed the incorporation of maximum temperature and VPD information that are highly variable between regions. The combined benefits of these methods are reflected in an improved ability to describe yield in a comprehensive data base spanning wide range of soil and climate. We quantified the increase in frequency of the more stressful type of environments with realised climate change, particularly in Southern Queensland and Northern New South Wales. We also demonstrated how earlier flowering partially mitigates heat and drought stress although, at the expense of reduced crop growth duration and yield potential, and increased risk of chilling and frost stress, which are currently minor.

## Acknowledgments

This research was enabled by Grains Research and Development Corporation (GRDC) through the provision of data resources generated as part of the NVT program and provided through the NVT Resource Sharing pathway. We also thank GRDC for investment in project UOQ2402-010RTX ‘Fast tracking deployment of chickpea heat tolerance to develop chickpea varieties with improved high temperature tolerance’. We also thank Chickpea Breeding Australia for sharing of data resources. Acknowledgment is made to the APSIM Initiative which takes responsibility for quality assurance and a structured innovation programme for APSIM’s modelling software, which is provided free for research and development use (see www.apsim.info for details).

---

**Supplementary Figure 1.**
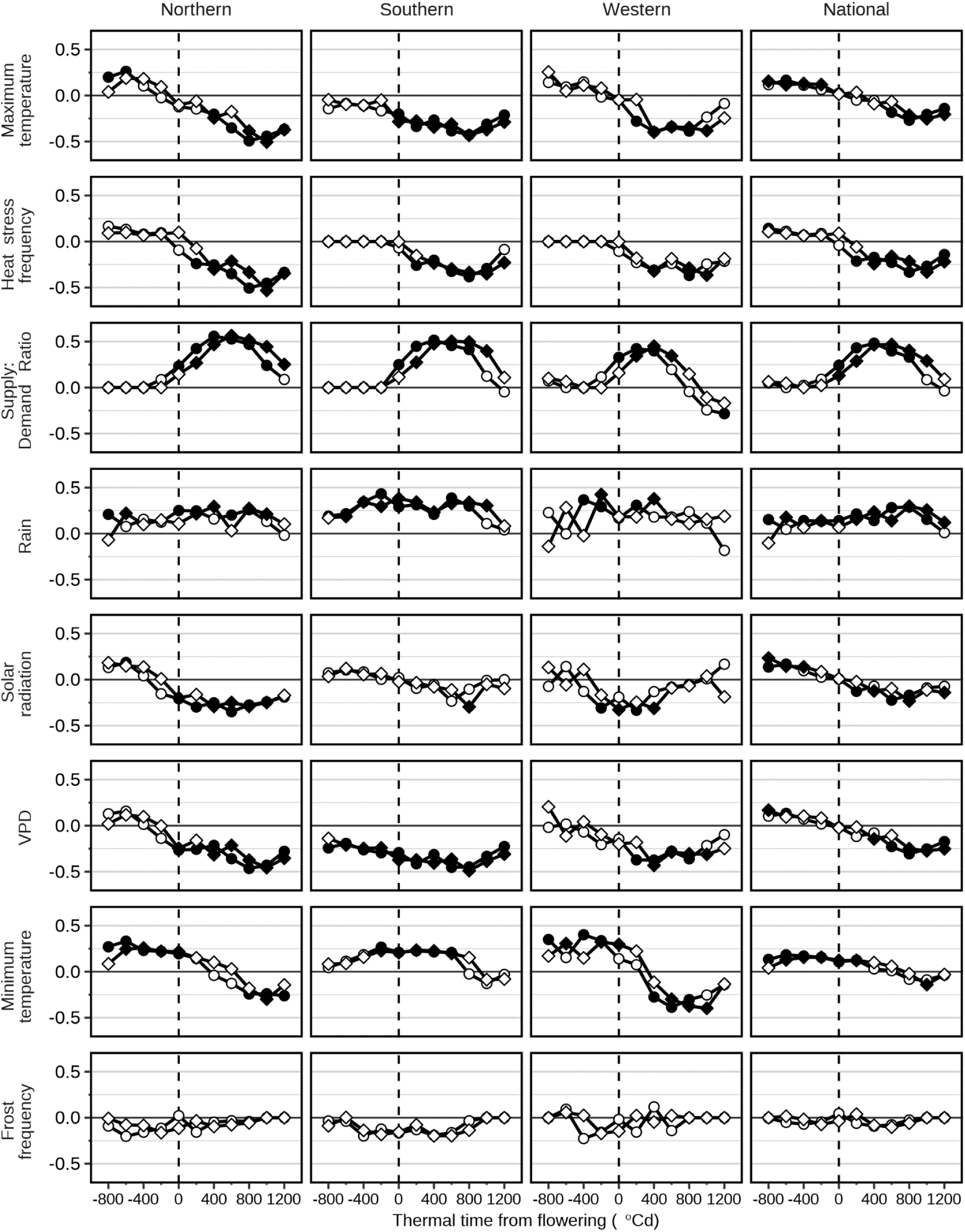
Pearson’s coefficient of correlation between Nataional Variety Trial mean yield and eight environmental factors. Factors are centred on the average simulated flowering date of variety Sonali (diamnonds) and PBA HatTrick (circles), and averaged every 200 °Cd. Correlations presented at the regional and continental scales. Water supply:demand also simulated with PBA HatTrick. Symbols indicate Benjamini-Hochberg adjusted *p* ≤ 0.05 (closed) and *p* > 0.05 (open). We defined thresholds as heat stress, is maximum temperature > 30°C, and frost, i.e., minimum temperature < 0 °C. VPD is vapour pressure deficit.

**Supplementary Figure 2.**
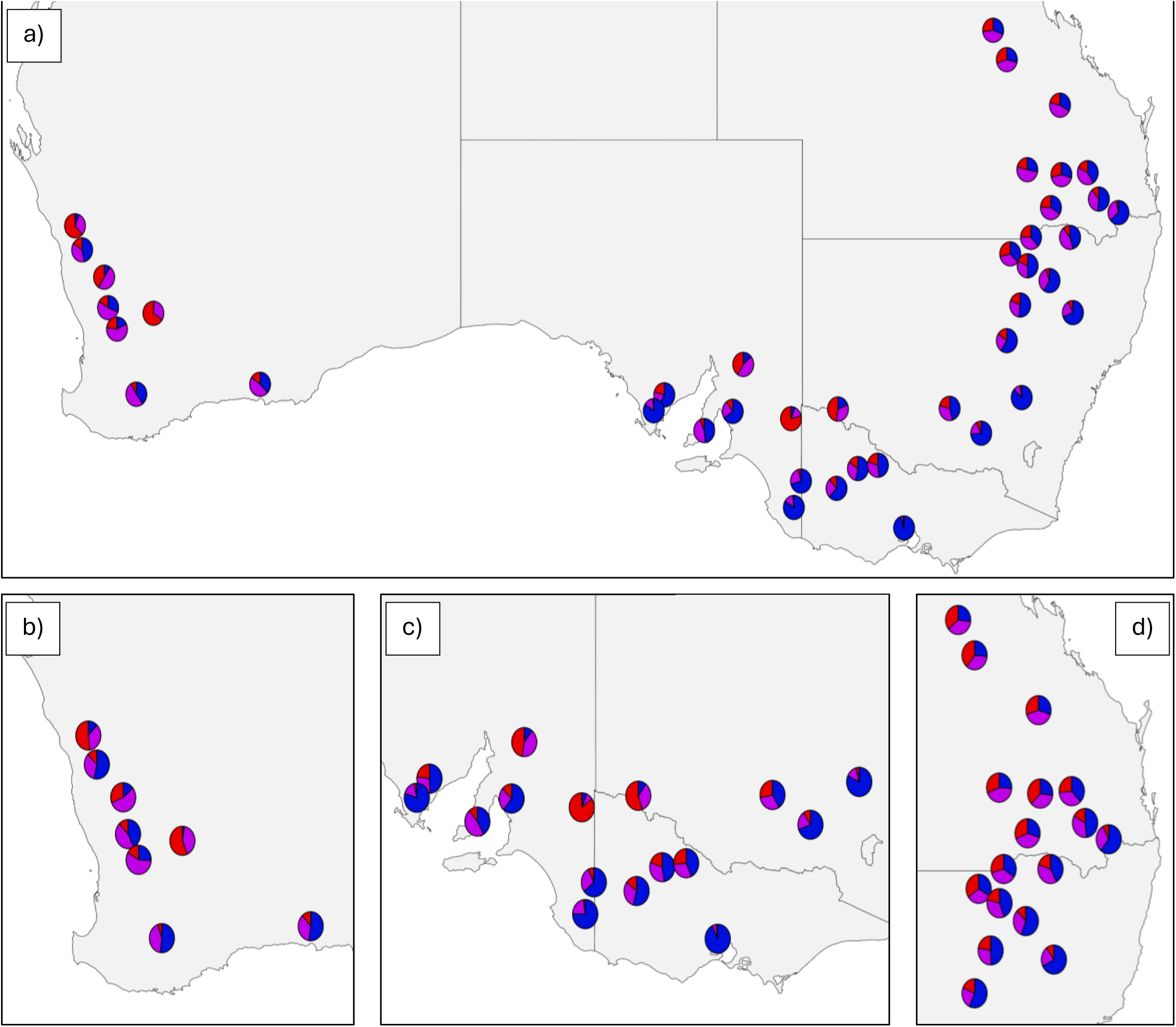
Spatial variation in the frequency of water stress (supply:demand) for a) environment types identified for the whole continent and environment types identified in each of the b) western, c) southern and d) northern regions. Nearby locations with similar frequency profiles were grouped to minimise overlap between pie charts. Based on simulations from 1957 to 2023. The characteristics of ETs at the regional and continental scales are presented in Fig. 5.

