## Supplemental Table 1 and 2 for "Spatial probabilistic patterns of joint heat and water stress of chickpea in Australia"

**Supplementary Table 1.** Correlation coefficients for the average water supply:demand ratio (SDR), maximum temperature (MaxT), minimum temperature (MinT), solar radiation (Radn) and vapour pressure deficit (VPD) from 200 ^o^Cd to 600 ^o^Cd after flowering across the southern region.

|  | Supply:demand | Maximum Temperature | Minimum Temperature | Radiation |
| --- | --- | --- | --- | --- |
| Maximum Temperature | -0.57 |  |  |  |
| Minimum Temperature | -0.13 | 0.40 |  |  |
| Radiation | -0.46 | 0.73 | 0.08 |  |
| VPD | -0.60 | 0.96 | 0.13 | 0.76 |

**Supplementary Table 2.** Correlation coefficients for the average water supply:demand ratio (SDR), maximum temperature (MaxT), minimum temperature (MinT), solar radiation (Radn) and vapour pressure deficit (VPD) from 200 ^o^Cd to 600 ^o^Cd after flowering across the northern region.

|  | Supply:demand | Maximum Temperature | Minimum Temperature | Radiation |
| --- | --- | --- | --- | --- |
| Maximum Temperature | -0.52 |  |  |  |
| Minimum Temperature | -0.18 | 0.55 |  |  |
| Radiation | -0.36 | 0.57 | 0.02 |  |
| VPD | -0.54 | 0.93 | 0.22 | 0.66 |

**Supplementary Table 3.** Correlation coefficients for the average water supply:demand ratio (SDR), maximum temperature (MaxT), minimum temperature (MinT), solar radiation (Radn) and vapour pressure deficit (VPD) from 200 ^o^Cd to 600 ^o^Cd after flowering across the western region.

|  | Supply:demand | Maximum Temperature | Minimum Temperature | Radiation |
| --- | --- | --- | --- | --- |
| Maximum Temperature | -0.73 |  |  |  |
| Minimum Temperature | -0.28 | 0.58 |  |  |
| Radiation | -0.68 | 0.68 | 0.16 |  |
| VPD | -0.76 | 0.96 | 0.35 | 0.74 |
